# Selecting coral species for reef restoration

**DOI:** 10.1101/2021.11.03.467181

**Authors:** Joshua S. Madin, Michael McWilliam, Kate Quigley, Line K. Bay, David Bellwood, Christopher Doropoulos, Leanne Fernandes, Peter Harrison, Andrew S. Hoey, Peter J. Mumby, Juan C. Ortiz, Zoe T. Richards, Cynthia Riginos, Nina Schiettekatte, David J. Suggett, Madeleine J. H. van Oppen

## Abstract

1. Humans have long sought to restore species, but little attention has been directed at how to best select a subset of foundation species for maintaining rich assemblages that support ecosystems, like coral reefs and rainforests that are increasingly threatened by environmental change.
2. We propose a two-part hedging approach that selects optimized sets of species for restoration. The first part acknowledges that biodiversity supports ecosystem functions and services, and so it takes precaution against loss by ensuring an even spread of phenotypic traits. The second part maximizes species and ecosystem persistence by weighting species based on characteristics that are known to improve ecological persistence—e.g., abundance, species range and tolerance to environmental change.
3. Using existing phenotypic trait and ecological characteristic data for reef building corals, we identified sets of ecologically persistent species by examining marginal returns in occupancy of phenotypic trait space. We compared optimal sets of species with those from the world’s southern-most coral reef which naturally harbors low coral diversity to show these occupy much of the trait space. Comparison with an existing coral restoration program indicated that current corals used for restoration only cover part of the desired trait space and may be improved by including species with different traits.
4. *Synthesis and applications*. While there are many possible criteria for selecting species for restoration, the approach proposed here addresses the need to insure against unpredictable losses of ecosystem services by focusing on a wide range of phenotypic traits and ecological characteristics. Furthermore, the flexibility of the approach enables the functional goals of restoration to vary depending on environmental context, stakeholder values, and the spatial and temporal scales at which meaningful impacts can be achieved.

## 1. Introduction

The rate and extent of environmental change experienced by contemporary ecosystems have resulted in major deviations from their historical state (Hobbs et al., 2011). Protection alone may no longer suffice to preserve biodiversity and related ecosystem functions and services, and active restoration is increasingly considered essential. The objective of ecosystem restoration is, through human intervention, to recover a disturbed or degraded ecosystem as far as possible towards some preferred previous state. Interventions can be direct, such as propagation and field deployment of habitat builders through seeds (Orth et al., 2020), propagules (Vanderklift et al., 2020), early recruits (Fredriksen et al., 2020) or parts of adult tissues (Rinkevich, 2014; Page et al., 2018). Indirect interventions, such as physical stabilization of degraded reef structures and removal or macroalgae are also possible (Ceccarelli et al., 2020).

To date, the augmentation or reintroduction of one or few species has been the most common approach, such as the restoration of the endangered Caribbean coral species *Acropora cervicornis* and *A. palmata* (Ladd et al., 2019), the reintroduction of the gray wolf across parts of Europe and North America (Ripple et al., 2014), and assisted colonization of the Tasmanian Devil to the Australian mainland (Brainard, 2020). However, climate change is now affecting many assemblages of foundation species in most if not all the world’s ecosystems, including forests, kelp beds, and coral reefs, leading to a necessary broadening of focus of restoration activities to encompass more species and their broader contributions to ecosystem functioning (Laughlin, 2014; Coleman &Bragg, 2020).

## 2. Selecting species for restoration

Prioritizing sets of species for the restoration of biodiverse ecosystems is a challenging task. Some approaches focus on the roles that species play in providing particular ecosystem goods or services, including carbon storage in rainforests (Strassburg et al., 2020) or reef accretion on coral reefs for coastal protection (Bellwood et al., 2019). Another common focus is on keystone species: species that maintain the organization, and stability of their communities, and have disproportionately large, inimitable impacts on their ecosystems (Hale &Koprowski, 2018). Alternatively, weedy pioneer species may quickly restore habitat functions such as providing shelter or stabilizing substratum; this is exemplified by the emphasis on fast growing acroporids in coral gardening and larval-based restoration initiatives (Bostrom-Einarsson et al., 2020). However, approaches for species selection that consider multiple ecological, functional and logistical criteria are rare (Suding et al., 2004; Lamb, 2018). Some examples exist for forest restoration (Meli et al., 2013), and some have used linkages between phenotypic traits and ecosystem services to select species (Giannini et al., 2017). Given the rapid growth in ecological and phenotypic trait databases across taxa globally (Gallagher et al., 2020), we propose a different approach that maximizes success in the context of a changing environment and uncertain future; we contend that hedging approaches should become an integral part of selecting sets of species for restoration.

## 3. Anticipating future ecosystems

An overarching challenge is that restoration initiatives need to anticipate future ecosystem states which are expected to be very different due to the escalating impacts of climate change (Rogers et al., 2015; Gaitán-Espitia &Hobday, 2021). Faced with complex ecosystems, multiple threats to biodiversity and limited funding, conservation practitioners must prioritize investment into different management options, including restoration actions, and difficult decisions must be made about which sets of species to allocate resources (Game et al., 2018). Strategic decisions must be made about supporting sets most likely to do better to improve future persistence and resilience, or helping sets that will struggle through a period of elevated and prolonged stress; especially those that are already closer to their existing physiological limits, like reef-building corals (Coles &Brown, 2003). Protecting habitat-forming species such as corals is imperative for securing the ecosystem services they provide, such as reef building, habitat and food provisioning for commercially important species, coastal protection, and biomedical, social, cultural and recreational opportunities. Given the sheer number of interacting ecological and social considerations, as well as the massive diversity of taxa in ecosystems such as coral reefs, deciding which species to select for restoration is complex, and the answer can change with each new consideration. Hedging strategies reduce risk of future loss of functions and services, but this could come at the cost of the optimal future solution (Nee &May, 1997).

## 4. Two-part hedging approach

The restoration of ecosystems via foundation assemblages should not only consider which species are most likely to persist in the future, but also which species provide essential contributions to the current suite of ecosystem services. To balance these considerations, we argue for a two-part process when selecting sets of foundation species. The first part centers on the maintenance of certain ecosystem services provided by foundation species, ranging from habitat engineering to phylogenetic diversity. However, rather than targeting specific properties that support ecosystem services, we propose to minimize their loss by maximizing phenotypic variation (e.g., selecting species that occupy trait space evenly). To demonstrate, we focus on reef-building corals and used the dataset from McWilliam et al. (2018) containing seven phenotypic traits (growth rate, corallite width, rugosity/branch spacing, surface area per unit volume, colony height, maximum colony size/diameter, and skeletal density; Fig. 1A) for the 396 reef coral species from Eastern Australia. These trait data enabled us to capture important dimensions of species life history globally, ranging from fast to slow growth (Darling et al., 2012), fragile to robust morphologies (Zawada et al., 2019), and small to large colonies that drives up colony fecundity (Alvarez-Noriega et al., 2016).

**Figure 1.**
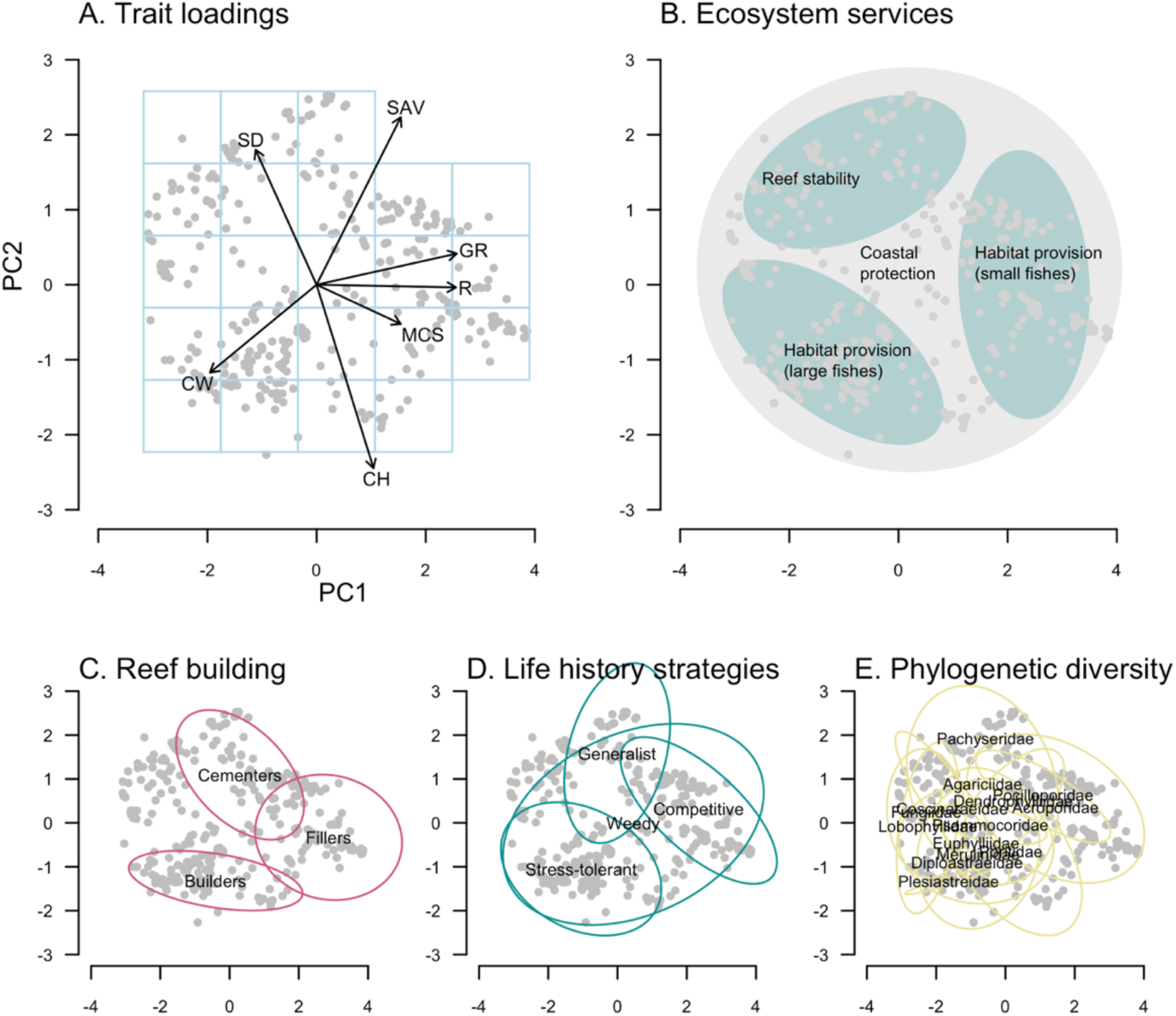
Phenotypic trait space as represented by the first two principal component axes for 396 Eastern Australia coral species. (**A**) The 5 by 5 cell grid used to calculate occupancy (see Supplementary Methods) and the trait loadings: growth rate (GR), corallite width (CW), rugosity/branch spacing (R), surface area per unit volume (SAV), colony height (CH), maximum colony size/diameter (MCS), and skeletal density (SD). (**B**) A schematic showing that some ecosystem services require species broadly across the space (e.g., reef-building and coastal protection) whereas others only a limited region of the space (e.g., reef stability and habitat provision). The schematic is illustrative only; not based on data. (**C**) Goreau’s (1963) categories of essential reef builders. (**D**) Darling et al.’s (2012) life history categories. (**E**) Veron’s (2000) families. Ellipses in **C**-**E** are 95% confidence for each category’s centroid.

Our reasons for maximizing phenotypic variation are four-fold. First, mechanistic linkages between phenotypic traits of species with the functions and services they provide are poorly understood, especially in marine systems (Bellwood et al., 2019). Therefore, it is difficult to ascertain regions of trait space responsible for particular services. Second, maximizing phenotypic variation increases life history variation, and therefore minimizes the risk of wholesale species loss (i.e., response diversity), because no species is at a selective optimum in all situations and environments (Stearns, 1992). Third, while some services may occupy small regions of trait space, maintaining multiple ecosystem services requires different groups of species covering the entire trait space (Fig. 1B). For example, Goreau’s (1963) reef building groups—builders, fillers and cementers—span most of coral phenotypic trait space (Fig. 1C; see Supplementary Methods); as do Darling et al.’s (2012) life history groups (Fig. 1D). Finally, hedging strategies also act to increase phylogenetic diversity because many life history traits are phylogenetically conserved and so occupy limited regions of trait space (Fig. 1E) (Westoby et al., 2002).

The second part of the process is to select species based on ecological characteristics that make them better equipped to avoid depletion and resist or recover from large-scale events, such as marine heatwaves. For example, species with small range sizes and small local populations generally have a higher extinction risk (Staude et al., 2020), while species with higher local abundances and larger range sizes tend to bounce back faster following disturbance (Halford et al., 2004). Meanwhile, some species are more tolerant to disturbances and changes that are expected to become more frequent or more intense in the future through rapid adaptation. To demonstrate, we used three ecological characteristics from the Coral Trait Database (Madin et al., 2016; ecological abundance, geographic range size, and thermal bleaching susceptibility) to estimate emergent characteristics of populations that are likely to make them more persistent.

## 5. Eastern Australia case study

While there are many definitions of phenotypic trait diversity (Villéger et al., 2008), our goal under a hedging strategy was to evenly capture the largest area of trait space with the fewest species in a biogeographic region, therefore ensuring a spread of species along important, often orthogonal, trait dimensions. This goal was accomplished by iteratively removing the species closest to other species in multi-dimensional trait space until some prerequisite number of species remained (see Supplementary Methods). Ecological persistence was integrated by multiplying species distances in trait space by standardized ecological characteristics during each iteration (i.e., species with standardized values closer to 1—i.e., large ranges, ecologically common, and resistant to thermal bleaching—were considered ecologically persistent and so less likely to be removed during an iteration; Supplementary Methods). Occupancy of trait space was measured using a 5 by 5 grid of cells (Fig. 1A). The results are reported in Fig. 2A.

**Figure 2.**
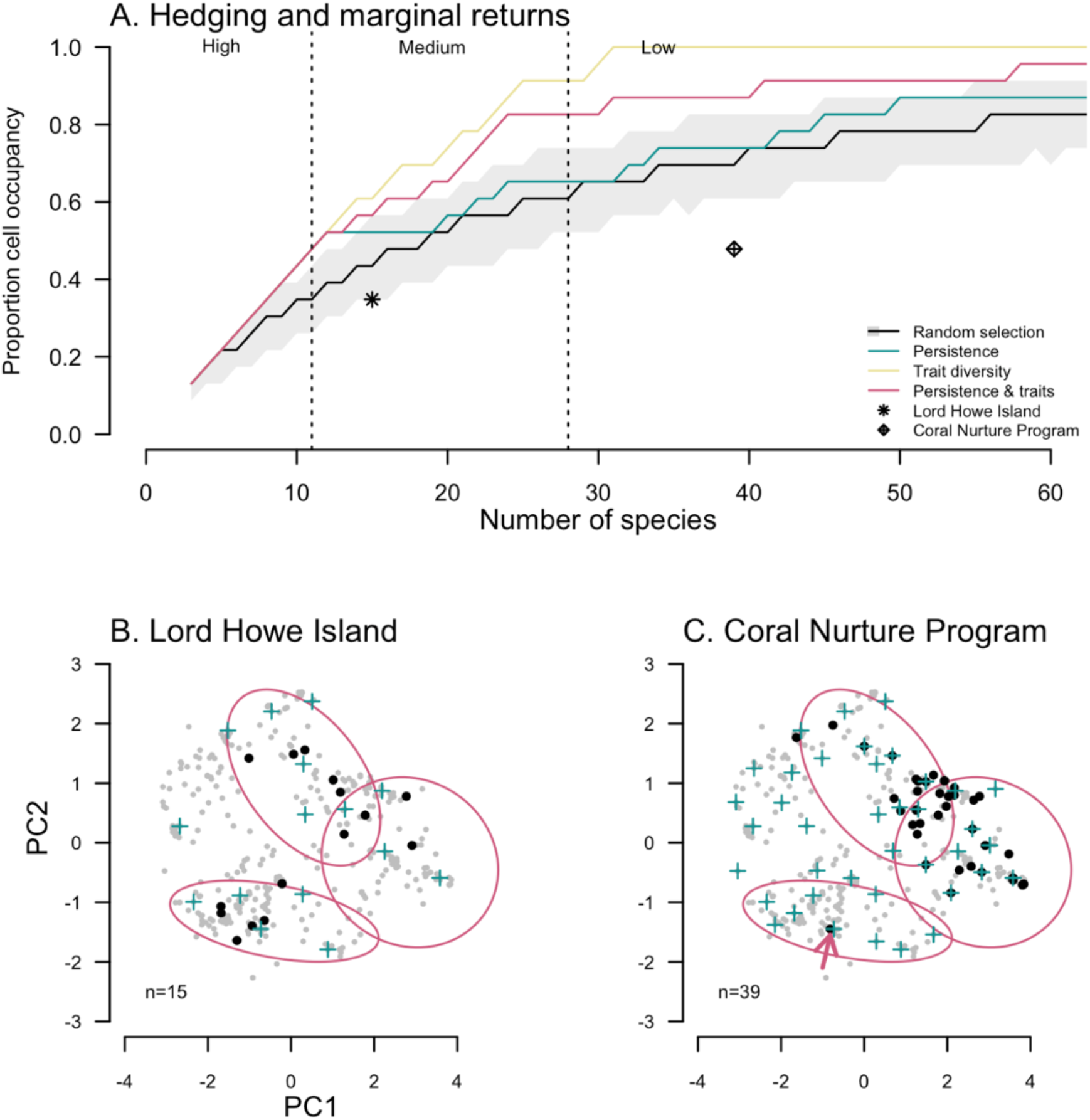
(**A**) Proportion occupancy of selected Eastern Australia species in trait space as a function of number of species using a 5 by 5 cell grid (see Fig. 1A). Regions of high, medium and low marginal returns delineated with dotted vertical lines. Included are symbols that show trait diversity and redundancy for common species at the world’s southernmost coral reef, Lord Howe Island (*n*=15), and for the Coral Nurture Program (*n*=39). The grey shaded region shows 95% CIs for randomized species selection. (**B**) Lord Howe Island (*n*=15) and (**C**) Coral Nurture Program (*n*=39) species are shown in trait space as black points. Red ellipse are Goreau’s (1964) reef building categories (centroid 95% confidence regions). Blue crosses are species selected by the two-part hedging process for 15 and 39 species, respectively. Red arrow in **C** is *Galaxea fascicularis*.

The hedging approach highlighted several important points for selecting species (Fig. 2A).

- Selecting species that maximize distances in trait space resulted in superior levels of occupancy (Fig. 2A, yellow curve). However, selecting species randomly yields relatively high levels of occupancy (see also Nee and May, 1997), and so might be used when phenotypic trait data are missing or incomplete (Fig. 2A, black curve and grey 95% confidence band).
- Selecting species based on ecological persistence alone (i.e., hedging against future loss of restored species) was broadly equivalent to random selection with regard to trait space occupancy (Fig. 2A, green curve). This result suggests that species likely to weather future conditions are distributed randomly across the trait space, which is a positive outcome.
- Integrating ecological persistence with trait diversity marginally reduces trait space occupancy compared with using traits diversity only, because species occupying distinct regions of trait space are being removed because they are unlikely to survive future conditions. That is, this two-part hedging approach protects against selecting high risk species in the effort to hedge against ecosystem service losses.
- In terms of how many species to select, marginal returns in trait space occupancy as species are added is high (approximately 5% occupancy per species) up until approximately 11 species; medium (approximately 2% per species) between from 11 to 28 species; and low (<0.3% per species) above *n*=28 (Fig. 2A, vertical dotted lines). The general patterns shown in Fig. 2A were robust to the cell size of the grid used to calculate occupancy (Fig. S2); with the proviso that more species are required to maintain specified levels of occupancy for finer grids. Our hedging approach therefore clearly quantifies “bang for buck”: that is, the amount of future hedging per species to be restored.
- We compared our hedging results with those of the southern-most accreting reef in eastern Australia, Lord Howe Island, and a coral restoration program, the Coral Nurture Program. The 15 common coral species found at Lord Howe Island occupy trait space no differently to randomly selecting species (Fig. 2A, asterisk) and occupy broad regions that likely support reef building (Fig. 2B, red ellipses). Coral Nurture Program, which selected 39 species primarily based on commonness and ease of out-planting (i.e., mostly branching species), occupies only 40-50% of the possible trait space (Fig. 2A, diamond) and does not occupy regions of trait space with reef “builders” (with the exception of one species, *Galaxea fascicularis*, indicated by the red arrow, Fig. 2C). We identify in both examples the species that would have been selected from the total community of corals found based on the two-part hedging process (Fig 2B, C, blue crosses).
- Finally, the two-part approach can be flexible and expanded upon. By way of example, we use the hedging approach to select 20 Eastern Australia species (medium marginal returns in Fig. 2A) based on two management scenarios: first, selecting species most likely to persist in the future based on high scores for ecological abundance, geographic range size and bleaching resistance (Fig. 3A) and, second, selecting those most vulnerable to thermal bleaching, but with high scores for ecological abundance and geographic range size again (Fig. 3B). The first columns in each panel are sets of species based on ecological characteristics alone. The second columns are sets of species when additionally considering trait space occupancy. The third columns are sets of species when expanding the approach to also considering a species’ ease of restoration (here we rank morphologies by ease of restoration as per Boström-Einarsson et al., 2020; see Supplementary Methods). Fig. 3 shows the impact of switching species that are better ecologically, but are difficult to restore based on our criteria, while simultaneously retaining an even spread of species in the trait space.

**Figure 3.**
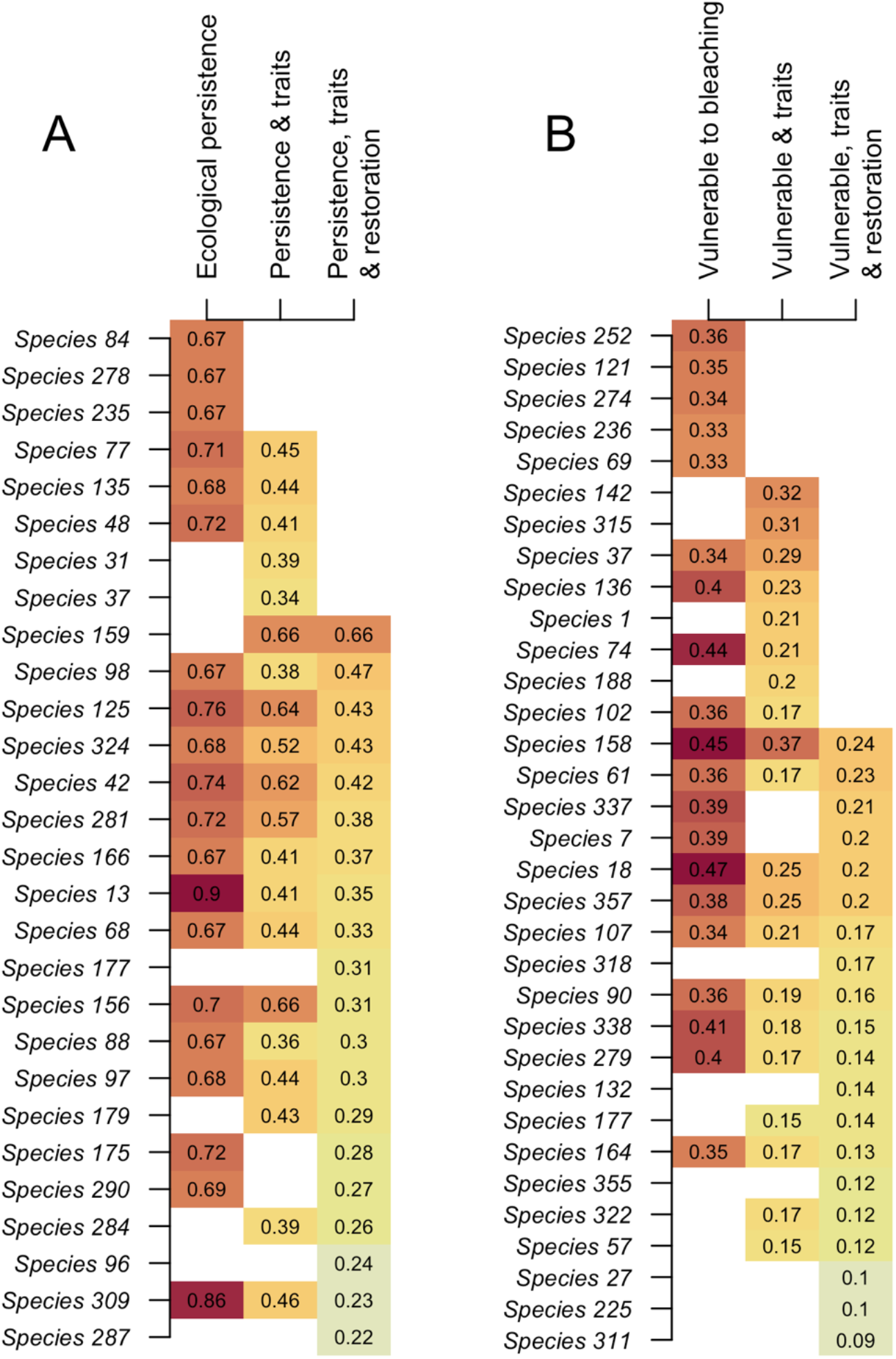
Sets of 20 species for different stages of the hedging process when focusing on (**A**) ecologically persistent species in terms of range size, local abundance and bleaching resistance, and (**B**) ecologically persistent species in terms of range size and local abundance, but that are vulnerable to bleaching. First columns consider ecological characteristics only. Second columns consider ecology and trait space occupancy. Third columns considering a new variable: a species ease of restoration based on morphology (see Supplementary Methods). Values (and heat colors) range between 0 and 1 correspond with a species normalized selection score at successive stages. Species names have been anonymized to avoid overinterpretation of the results.

## 6. Conclusions

The two-part hedging approach outlined here, and demonstrated with reef-building corals, identified species for active restoration projects based on both diversity of life history trait values and ecologically beneficial characteristics. Selection based on ecological characteristics are important for hedging against future species loss, while trait diversity is important for hedging against the loss of certain ecosystem services, reef-building groups, life history categories, and phylogenetic diversity. The importance of individual species for ecosystem processes, functions and services are poorly understood, particularly for reef-building corals. Furthermore, selecting species based on ecosystem services only is likely to vary across systems depending on the specific environment, and the values of the local stakeholders (Bellwood et al, 2019). Our tactic was therefore to prioritize an even spread of species across the trait space rather than prioritizing particular phenotypic trait values or targeting regions in the trait space (Fig. 1). When selecting species for restoration, a species-oriented focus may favor rare or depleted species with low persistence. However, ecosystem restoration based on trait diversity is more likely to be robust if it favors foundation species that are dominant and persistent (as we advocate here), and therefore more likely to maintain a broad range of ecosystem services. The number of species that can be selected for a restoration project will ultimately depend upon project goals, as well as resource and logistical constraints; however, marginal returns in trait space occupancy can help managers and practitioners decide where to draw the best line. The flexible species selection process developed here can serve as a framework for such decisions, and can serve an important role even as the goals of restoration are refined based on an improved knowledge of ecosystem services, successful restoration methods, diverse stakeholder values, and the scales at which restoration is most effective.

## Authors contributions

All authors conceived the idea during a working group meeting organized by MJHvO and KQ. JSM and MM developed the idea, gathered data and ran analyses. JSM, MM, KQ and MJHvO wrote the first draft. All authors critically revised drafts and added intellectual content.

## Acknowledgements

The workshop was funded by the Australian Research Council Laureate Fellowship FL180100036 to MJHvO. MJHvO, CD and LKB acknowledge the Reef Restoration and Adaptation program, which is funded by the partnership between the Australian Governments Reef Trust and the Great Barrier Reef Foundation. We also acknowledge the National Science Foundation (1948946 to JSM) and Australian Research Council (LP160101508 to ZR and FL190100062 to DRB).

## Data availability statement

All data and code are available at https://github.com/jmadinlab/species_choice

