## Supplementary Methods for "Selecting coral species for reef restoration"

Joshua S. Madin[1]*, Michael McWilliam[1], Kate Quigley[2], Line K. Bay[3], David Bellwood[4], Christopher Doropoulos[5], Leanne Fernandes[6], Peter Harrison[7], Andrew S. Hoey[3], Peter J. Mumby[8], Juan C. Ortiz[2], Zoe T. Richards[9], Cynthia Riginos[8], Nina Schiettekatte[1], David J. Suggett[10], Madeleine J. H. van Oppen[3,11]

1. Hawai‘i Institute of Marine Biology, University of Hawai‘i at Manoa, Kāne‘ohe, Hawai‘i, USA
2. Minderoo Foundation, Perth, WA, Australia
3. Australian Institute of Marine Science, Townsville, Queensland, Australia
4. College of Science and Engineering, James Cook University, Townsville, Queensland, Australia
5. CSIRO Oceans & Atmosphere, Brisbane, Queensland, Australia
6. Great Barrier Reef Marine Park Authority, Townsville, Queensland, Australia
7. Marine Ecology Research Centre at Southern Cross University, New South Wales, Australia
8. School of Biological Sciences, The University of Queensland, St. Lucia, Queensland, Australia
9. Coral Conservation and Research Group, Trace and Environmental DNA Laboratory, School of Molecular and Life Sciences, Curtin University, Bentley, Western Australia, Australia
10. University of Technology Sydney, Climate Change Cluster, Sydney, New South Wales, Australia
11. School of BioSciences, The University of Melbourne, Parkville, Victoria, Australia

*

### Supplementary Methods

*Functional trait space*

For our demonstration, we use a dataset for 396 species found along the east coast of Australia from McWilliam et al. (2018) with the following traits: growth rate, corallite width, rugosity/branch spacing, surface area per unit volume, colony height, maximum colony size/diameter, and skeletal density. The trait data enabled us to capture important dimensions of species life history, ranging from fast to slow growth (McWilliam et al., 2022), fragile to robust morphologies (Zawada et al., 2019), and small to large colonies that drives up colony fecundity (Alvarez-Noriega et al., 2016). The trait space presented main text in Fig. 1 was calculated using a principal components analysis (PCA) of the seven traits. We used the first two PC axes which captured approximately 70% total trait variation.

We viewed ecosystem values across the trait space through several analyses. Fig. 1B was a general qualitative schematic for a selection of processes, functions and services. For reef building (Fig. 1C), we adopted Goreau's (1963) classification of reef species into builders, fillers and cementers; an approach that has been supported by modern synthesis (González-Barrios & Alverez Filip 2018). Within the McWilliam et al. (2018) trait space, builders were classified as species with the highest values of size, height, and volume; fillers with largest values of size and rugosity; and cementers with the largest sizes and smallest height values. For life histories (Fig. 1D), we adopted Darling et al.'s (2012) categories, which were extracted from the Coral Trait Database. For phylogeny (Fig. 1E), we used Veron's (2000) taxonomic families. Groups were presented in trait space as the 95% confidence ellipses for centroids.

*Phenotypic diversity*

While there are many definitions of phenotypic trait diversity (Villéger et al., 2008), our goal under a hedging strategy was to evenly capture the largest area of trait space with the fewest species, and therefore to ensure a spread of species along important trait dimensions. This goal was accomplished in several ways. The first was to iteratively removing the species closest to other species in the two-dimensional area defined by PC1 and PC2 until a given number of species *n* remained. We used the nearest neighbor distances using the *nndist* function in the *spatstat* package and only considering the single closest species (i.e., *k*=1; Baddeley et al., 2015). The second approach was to iteratively removing species with the smallest Voronoi cell areas in the two-dimensional area defined by PC1 and PC2 using the *voronoi.mosaic* function in the *tripack* package (Renka et al., 2020). The third was to use the *hypervolume* package (Blonder et al. 2022) and quantifying each species contribution to the total volume using the *kernel.contribution* function (Mammola & Cardoso 2020). Nearest neighbor distances, Voronoi areas and hypervolume contributions were normalized at each iteration by dividing by the maximum of the respective values. The three methods are presented in Fig. S1 for groups of 20 species. The Voronoi area method tended to bias species at the periphery of trait space for which areas could not be calculated. While the hypervolume approach provides the most robust approach, because it operates over all axes of variation, calculating hypervolumes was computer intensive and not practical for testing multiple scenarios for 396 species. Therefore, we present the nearest neighbor method in our perspective to illustrate the hedging process. However, the best method or combination of methods to be used should be the basis of further study.

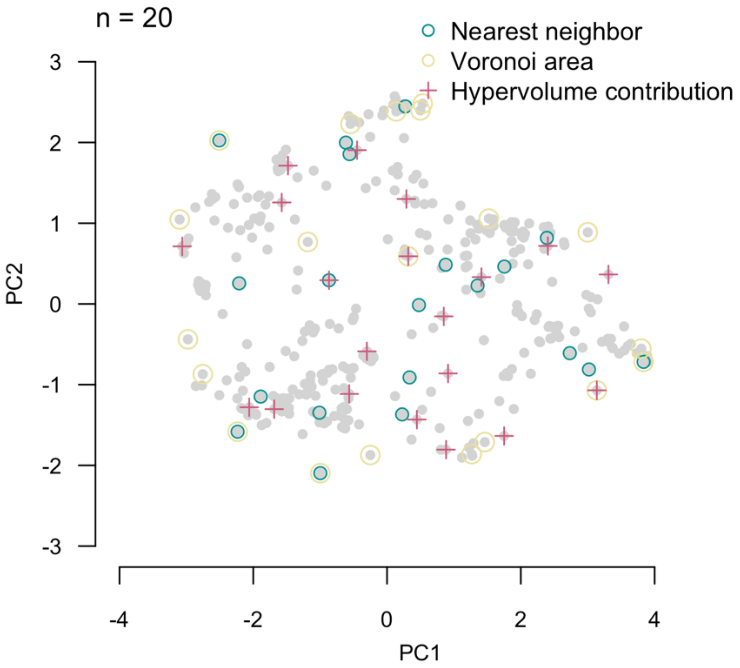


**Fig. S1**. Selecting set of 20 species in trait space using three methods. The nearest neighbor and Voronoi area methods operate on the first two PC axes; whereas the hypervolume contribution operates on the 7-trait-dimension hypervolume, but is presented here in PC space.

We used a grid-based approach to assess trait diversity, whereby a grid of a given resolution was superimposed onto the trait space (e.g., a 5 by 5 cell grid is shown in Fig. 1A and a 10 by 10 cell grid in Fig. S2). Trait diversity was the proportion of possible grid cells with at least one species; redundancy was the mean number of species in occupied possible grid cells.

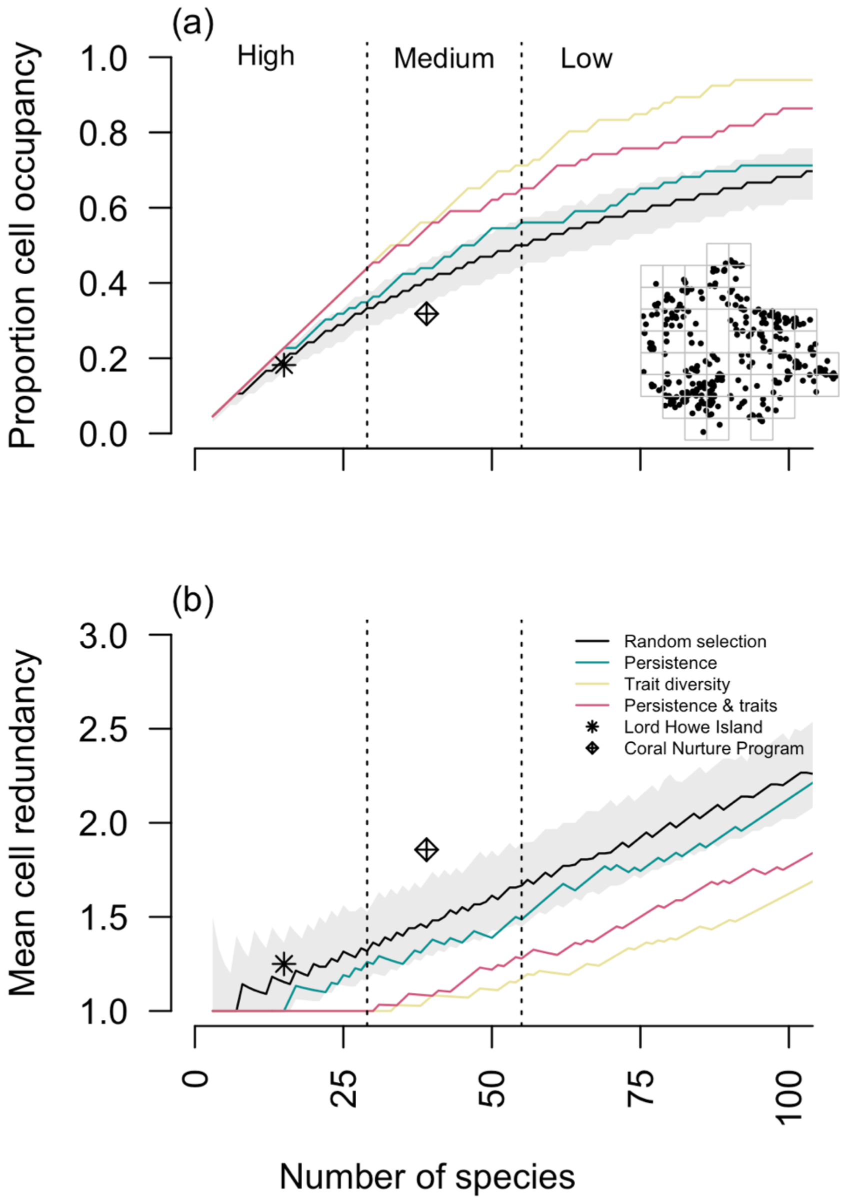


**Figure S2**. (**a**) Proportion occupancy and (**b**) redundancy in trait space as a function of number of species n using a 10 by 10 cell grid (inset in panel **a**). Regions of high, medium and low marginal returns delineated with dotted vertical lines. The asterisks show trait diversity and redundancy for the 15 species found at Lord Howe Island. The grey shaded region shows 95% CIs for randomized species selection.

*Ecological persistence*

For the second part of the hedging process, we focused on three ecological characteristics (ecological abundance, geographic range size and thermal bleaching susceptibility) broadly defined as characteristics contributing to ecological persistence of reef building coral species. We acknowledge that these characteristics will depend on current taxonomic designations that are currently being revised (Cowman et al., 2020). We used typical abundance of species data from Veron (2000) and geographic distribution data from Hughes et al. (2013), both downloaded for the 396 species from the Coral Trait Database (Madin et al., 2016). Ecological abundance was categorized by Veron (2000) as common, uncommon and rare, which we normalized as 1, 0.5 and 0.25, respectively. Geographic extent was normalized by dividing the range size of each species by the maximum range size for a species. Normalizing puts characteristics on the same scale (i.e., between 0 and 1).

Thermal bleaching susceptibility is an increasingly relevant characteristic for coral ecological persistence, but it is context dependent, highly variable, and poorly understood. Nonetheless, we use the Coral Bleaching Index (BI) from Swain et al. (2016) to demonstrate how this variable might be included in the triage analysis. BI is a value between 0 and 100, where higher values correspond with more thermally vulnerable species. Therefore, we normalized BI by dividing by 100 and subtracting the result from 1 (i.e., species with values closer to 1 are more resistant to bleaching based on Swain et al. [2016]). BI values were available at the species level for 212 of the species, and therefore genus level BIs were used for the remainder of the analysis in order to retain all 396 species.

Restoration of reef corals is a relatively new field (Hein et al., 2021), and so there is little long-term knowledge of what makes species more or less amenable to the restoration process. A meta-analysis of coral restoration studies ranked the use of coral growth forms in restoration projects (Boström-

Einarsson et al., 2020). While this ranking likely reflects a historical focus on coral gardening (i.e., fragmentation) as well as specific situations, such as the demise of branching *Acropora* species in the Caribbean, we nonetheless utilize this ranking as an index of species amenability to restoration. Species growth form was downloaded from the Coral Trait Database and species were ranked from 1 to 6: columnar (1), tabular (2), encrusting (3), foliose (4), massive (5), and branching (6), which includes corymbose and digitate. This ranking was normalized by dividing values by six.

Correlations among ecological characteristics and PC axes are shown in Fig. S3.

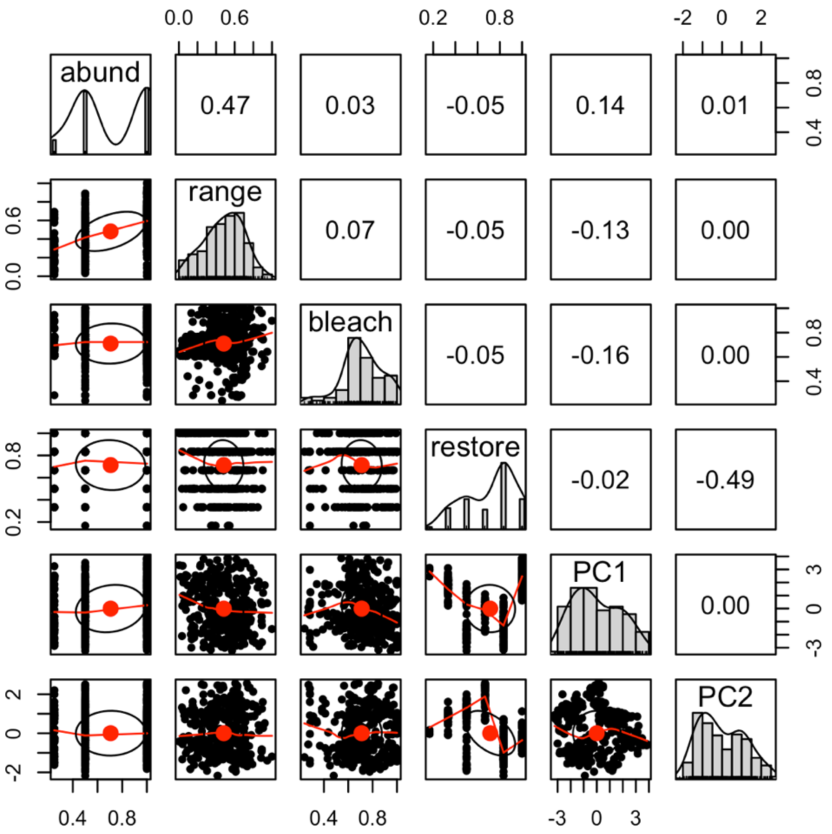


**Figure S3**. Pairwise associations between ecological characteristics, quantitative traits and principal components presented in the hedging analysis.

Weighting species by characteristics was done by multiplying normalized values together to get an overall persistence index. Species with values closer to 1—i.e., large ranges, common, and resistant to bleaching—were considered ecologically persistent. To redirect focus of species selection to vulnerable species, normalized variables were subtracted from 1 before proceeding. For example, we also explored the triage process for species that were wide ranging and common, but susceptible to bleaching (Fig. 3B).

*Iterative hedging process*

Sets of species were selected based on trait diversity or ecological persistence along, or as the integration of both. Fig 2A shows trait diversity—measured as occupancy of trait space—for a range of scenarios. The black line was occupancy when randomly selecting *n* species (grey shaded region showing 95% confidence for 1000 samples of *n* species). The blue line shows occupancy for *n* species selected based on ecological persistence alone. The yellow line shows occupancy for *n* species selected based on trait diversity alone. Finally, the red line show occupancy for *n* species selected by multiplying normalized distances in trait space by normalized ecological characteristics (i.e., abundance, range size and bleaching tolerance). So, this last scenario may occupy less of trait space, but the species are much more likely to persist.

Our final example was for a restoration program with the capacity to restore 20 species on a reef (Fig. 3), demonstrating how the set of species changes with restoration criteria. Fig. 3A focused on bleaching winners, and shows selected species when considering ecological persistence alone (first column), ecological persistence integrated with trait diversity (second column), and expanding to a new variable: a species amenability to restoration (third column). Fig. 3B focused on aiding bleaching losers, but follows the same process.
